# Intermediate induction of germline apoptosis maintains fertility and progeny fitness during temperature stress

**DOI:** 10.64898/2026.04.13.718325

**Authors:** Kristen A. Quaglia, Hannah N. Lorenzen, Samantha H. Oswald, Josephine M. Selvik, Lisa N. Petrella

## Abstract

Organisms must be able to maintain the ability to produce high quality offspring despite experiencing stressful conditions. It is unknown how *C. elegans* maintain the ability to produce offspring during moderate temperature stress just below the range of temperature that cause sterility. We evaluated apoptosis, fertility, and several progeny fitness metrics in no-apoptosis, high-apoptosis mutants, and in wild strains that varied in their fertility level during moderate temperature stress to understand if apoptosis is a strategy *C. elegans* use to maintain the ability to produce offspring during a moderate temperature stress. We found that apoptosis mutants were less fertile with less fit progeny compared to wild type under a moderate temperature stress. Wild strains isolated from the environment showed variability in the increase in apoptosis, levels of fertility, and measurements of progeny fitness observed. We also found that an intermediate induction of apoptosis trended with higher fertility and progeny fitness in wild strains under a moderate temperature tress. These results suggest that apoptosis within an optimal range in the *C. elegans* germline is a strategy used to maintain the ability to produce high quality offspring despite experiencing a moderate temperature stress. Many species also have germline apoptosis, so apoptosis may be a strategy other species use to maintain their own fertility when experiencing stress conditions

## Introduction

In the environment, organisms must have mechanisms to maintain their own fertility to produce fit progeny when they experience stress conditions. A variety of abiotic stressors, including increased temperature, can impact reproduction in a wide range of species (Salinas et al. 2006; Petrella 2014; Poullet et al. 2015; Fausett et al. 2021). Temperatures even a few degrees above the species optimal temperature can greatly reduce the fertility of organisms as diverse as mammals, flies, plants, and nematodes (Yaeram et al. 2006; Paupière et al. 2014; Petrella 2014; Poullet et al. 2015; Stazione et al. 2019; Fausett et al. 2021; Ci et al. 2025). The continued increase in surface temperatures due to climate change will affect the reproductive capacity of a breadth of organisms, making it imperative to identify mechanisms that contribute to the sustained production of healthy progeny during stress.

The nematode *Caenorhabditis elegans,* like other invertebrates, cannot control their own temperature, and thus are greatly impacted by any change in environmental temperatures. *C. elegans* have the highest fertility around 20°C and experience a decrease in fertility as temperatures increase from 20°C until they go sterile by 28°C (Petrella 2014; Poullet et al. 2015). *C. elegans* naturally occur across a large range of latitudes and thus experience a wide range of temperatures in their natural environments (Crombie et al. 2024), yet mechanisms *C. elegans* use to retain the ability to produce offspring despite experiencing these stress conditions remain largely unknown.

When experiencing the increasing environmental temperatures, the *C. elegans* germline has mechanisms to sustain fertility. In each U-shaped arm of the adult germline, oogenic nuclei start in the region distal to the uterus and move toward the bend in the germline (Hubbard and Greenstein 2005). During oogenesis, physiological apoptosis functions to remove approximately 50% of nuclei from the pool of cells that could become oocytes (Gumienny et al. 1999). Apoptosis occurs in the bend region of the germline where dying nuclei are engulfed into the surrounding somatic sheath cells, while their cytoplasmic components stream into cellularizing oocytes in the proximal arm of the germline (Gumienny et al. 1999; Wolke et al. 2007). These features of nuclear death but preserved cytoplasm have led the proposal that physiological apoptosis may function to remove damaged nuclei and also to supply cytoplasmic resources to the oogenic nuclei that remain in the germline (Bhalla and Dernburg 2005; Bailly and Gartner 2013). Previous studies have shown that germline apoptosis increases under a variety of stressors, including a moderate temperature stress (Salinas et al. 2006; Láscarez-Lagunas et al. 2014; Poullet et al. 2015; Compere et al. 2025). Thus, the increase in apoptosis observed under stress conditions could be cellular mechanism by which progeny fitness is insured either through inheritance of an intact genome and/or richer cytoplasmic components.

The canonical apoptosis pathway is highly conserved across a wide range of organisms. In *C. elegans,* the Bcl-2 homolog, CED-9, functions to repress apoptosis by sequestering CED-4, the Apaf-1 homolog, under non-apoptotic conditions (Ellis and Horvitz 1986). When apoptosis is induced, CED-9 function is inhibited, which releases CED-4 to activate the terminal caspase CED-3 (Hengartner et al. 1992). Dead nuclei are engulfed into the surrounding somatic sheath cells using CED-1, while most of the cytoplasm is shuttled into the remining nuclei through cytoplasmic streaming (Zhou et al. 2001; Wolke et al. 2007). Apoptosis is also kept at basal levels in the germ line through anti-apoptotic factors like GLA-3 and CPB-3 (Kritikou et al. 2006; Singh et al. 2017). GLA-3 and CPB-3 are both RNA-binding proteins that have been previously shown to inhibit germline apoptosis under normal conditions (Lettre et al. 2004; Kritikou et al. 2006; Singh et al. 2017). GLA-3 interacts with the MAP kinase MPK-1 in its role in apoptosis inhibition (Kritikou et al. 2006) and CPB-3 works through a largely unknown mechanism to inhibit apoptosis in the germline (Singh et al. 2017). The well characterized pro-and anti-apoptotic factors are advantageous for the study of the role of germline apoptosis in *C. elegans* in both unstressed and stressed conditions.

While most studies have focused on the N2 wild type while investigating stress response mechanisms, like moderate temperature stress, few have taken advantage of genetically distinct wild isolated strains. Previous studies that have used wild strains isolated from the environment have shown phenotypic and genotypic differences to the canonical laboratory wild type N2 (Reddy et al. 2009; Weber et al. 2010; Petrella 2014; Kwah and Jaramillo-Lambert 2023). Even within wild strains isolated from the environment, studies have identified phenotypic variation across strains, which can be used to understand the underlying evolution occurring in *C. elegans* (Petrella 2014; Zhang et al. 2021; Polk et al. 2025). The dynamic nature of organismal stress response mechanisms could indicate genetically distinct strains have a varied response to the same environmental stressor. Thus, it is likely that apoptosis, as a stress response mechanism in the germline, could also show phenotypic variation when studied in wild strains isolated from the environment. So, it is imperative to study these stress response mechanisms in a variety of wild strains.

Here we build upon previous works in other stressors to investigate progeny fitness metrics and apoptosis during moderate temperature stress in mutants that alter the level of apoptosis and a variety of wild strains more recently isolated from the environment. We show that an intermediate level of apoptosis in both unstressed and stressed conditions is most beneficial for fertility and progeny fitness. We also show that an intermediate induction of apoptosis is most beneficial for fertility and progeny fitness.

## Methods

### Strains and nematode culture

*C. elegans* were cultures under standard conditions (Brenner 1974) on NGM plates seeded with *E. coli* strain AMA1004 (Casadaban et al. 1983) at 20°C unless otherwise noted. Strains used for the experiments were MD701 *bcIs39 [lim-7p::ced-1::GFP + lin-15(+)]*, WS2170 *opIs110 [lim-7p::YFP::actin + unc-119(+)] IV,* MT2547 *ced-4(n1162) III*, MT1743 *ced-3(n718) IV*, CB3203 *ced-1(e1735) I*, WS2972 *gla-3(op216) I*, WS1137 *cpb-3(op234) I*, N2 (Bristol), ECA348, JU3135, ECA347, ECA705, JT11398, NIC166, EG4349, QG2813, QG2855, MY2212, NIC1107, LNP0002 *petIR2(V;bcIs39 [lim-7p::ced-1::GFP + lin-15(+)] N2>JT11398)*, LNP0003 *petIR3(V;bcIs39[lim-7p::ced-1::GFP + lin-15(+)] N2>N2)*, LNP0006 *petIR6(V;bcIs39[lim-7p::ced-1::GFP + lin-15(+)] N2>ECA348)*, LNP0033 *petIR14(V;bcIs39[lim-7p::ced-1::GFP+lin-15(+)] N2>MY2212)*, LNP0036 *petIR17(V;bcIs39[lim-7p::ced-1::GFP+lin-15(+)] N2>QG2855)*, LNP0047 *petIR20(V;bcIs39[lim-7p::ced-1::GFP+lin-15(+)] N2>NIC166)*, LNP0052 *petIR8(V;bcIs39[lim-7p::ced-1::GFP+lin-15(+)] N2>ECA347)*, LNP0055 *petIR11(V;bcIs39[lim-7p::ced-1::GFP+lin-15(+) N2>NIC1107),* LNP0069 *petIR25(V;bcIs39[lim-7p::ced-1::GFP+lin-15(+)] N2>EG4349)*, LNP0118 *petIR26(V;bcIs39[lim-7p::CED-1::GFP+lin-15(+)] N2>ECA705),* LNP0121 *petIR29(V;bcIs39[lim-7p::CED-1::GFP+lin-15(+)]) N2>JU3135),* LNP0124 *petIR32(V;bcIs39[lim-7p::CED-1::GFP+lin-15(+)]) N2>QG2813).* Some strains were provided by the CGC, which is funded by the NIH Office of Research Infrastructure Programs (P40 Od010440), and the *Caenorhabditis* Natural Diversity Resource (Crombie et al. 2024).

### Brood size assay

Worms were maintained at 20°C until the L4 stage, at which point L4 worms were cloned out onto individual NGM plates and placed at either 20°C, 26°C, or 27°C for the remainder of the experiment. Animals were allowed to lay eggs for 24 hours and were then transferred to a new NGM plate. Animals were transferred until no more eggs were laid. F1 progeny were counted 24 hours after P0 animals were transferred to the next plate. The brood size of P0 animals that died before the end of the reproductive period were excluded from analysis. Each strain was tested in at least three biological replicates per strain at each temperature for a total of n between 18 and 38 worms per strain per temperature. Data were analyzed using a Two-Way ANOVA with Tukey’s multiple comparisons using Prism 10 (Graphpad, San Diego, CA, USA).

### Embryo Survival assay

Worms were maintained at 20°C until the L4 stage, at which point L4 worms were transferred to an NGM plate and placed at either 20°C, 26°C, or 27°C for 24 hours. After 24 hours, P0 animals were cloned out onto individual NGM plates with thin bacterial lawns. Thin bacterial lawn NGM plates were made by spreading 50µl AMA1004 *E. coli* (Casadaban et al. 1983) in LB Broth onto blank NGM plates and allowed to dry overnight. After 6 hours, P0 animals were removed from plates and the number of embryos were counted. Plates were returned to experimental temperature for 24 hours. After 24 hours, the number of worms was counted to determine the percentage of surviving embryos. The percentage of embryo survival was calculated by the equation: (# worms / # embryos) * 100. Each strain was tested in at least three biological replicates for an n between 4 and 49 worms per strain per temperature. The data were analyzed by a Two-Way ANOVA with Tukey’s multiple comparisons using Prism 10 (Graphpad, San Diego, CA, USA).

### Acridine Orange Apoptosis

Worms were maintained at 20°C until the L4 stage, at which point L4 worms were transferred to an NGM plate and placed at either 20°C, 26°C, or 27°C for 24 hours. After 24 hours, young adult worms were placed onto NGM plates spotted with 20µl AMA1004 *E. coli* (Casadaban et al. 1983) overlayed with 50µg/ml acridine orange dye in 1XM9 and incubated in the dark for two hours at appropriate temperature. After two hours, worms were transferred to a fresh NGM plate with no dye for 10 min in dark, then transferred again to a fresh NGM plate for one hour and 50 min in dark. Individual worms were mounted on a 2% agarose pad in 10µM levamisole in 1X M9 without bacteria. Images were acquired using Leica Application Suite Advanced Fluorescence 3.2 software using Leica CTR6000 deconvolution inverted microscope with a Hamatsu Orca-R2 camera and Plan Apo 63x/1.4 numerical aperture oil objective. One gonad arm per animal was imaged. Images were taken in at least three biological replicates for an n between 15 and 17 worms per strain per temperature. Statistical analysis was performed by Two-Way ANOVA with Tukey’s multiple comparisons using Prism 10 (Graphpad, San Diego, CA, USA)

### ACT-5::YFP Apoptosis

Worms were maintained at 20°C until the L4 stage, at which point L4 worms were transferred to an NGM plate and placed at either 20°C, 26°C, or 27°C for 24 hours. After 24 hours, worms were mounted on a 2% agarose pad in 10µM levamisole in 1XM9 without bacteria. Images were acquired using Leica Application Suite Advanced Fluorescence 3.2 software using Leica CTR6000 deconvolution inverted microscope with a Hamatsu Orca-R2 camera and Plan Apo 63x/1.4 numerical aperture oil objective. One gonad arm per animal was imaged. Images were taken in at least three biological replicates for an n between 15 and 17 per genotype per temperature. Statistical analysis was performed using a Two-Way ANOVA with Tukey’s multiple comparisons using Prism 10 (GraphPad, San Diego, CA, USA).

### Creation of Near Isogenic Lines (NILs)

Near isogenic lines were created by crossing MD701 (*bcIs39 [lim-7p::ced-1::GFP + lin-15(+)])* hermaphrodites with wild isolate males. F1 male progeny were screened for presence of GFP and crossed to hermaphrodites of the same wild isolate background. Males from four subsequent generations were screened for GFP and crossed to hermaphrodites of the same wild isolate background. After five backcrosses, hermaphrodite F6s were screened for the presence of GFP. Hermaphrodites were allowed to make self-progeny until the GFP was homozygous.

### CED-1::GFP Apoptosis

Worms were maintained at 20°C until the L4 stage, at which point L4 worms were transferred to an NGM plate and placed at either 20°C, 26°C, or 27°C for 24 hours. After 24 hours, worms were mounted on a 2% agarose pad in 10µM levamisole in 1XM9 without bacteria. Images were acquired using Leica Application Suite Advanced Fluorescence 3.2 software using Leica CTR6000 deconvolution inverted microscope with a Hamatsu Orca-R2 camera and Plan Apo 63x/1.4 numerical aperture oil objective. One gonad arm per animal was imaged. Images were taken in at least three biological replicates for an n between 15 and 24 per strain per temperature. Statistical analysis was performed using a Two-Way ANOVA with Tukey’s multiple comparisons using Prism 10 (GraphPad, San Diego, CA, USA).

### Oocyte counting and measuring

Worms were maintained at 20°C until the L4 stage, at which point L4 worms were transferred to an NGM plate and placed at either 20°C, 26°C, or 27°C for 24 hours. For imaging, worms were mounted on slides with 2% agarose pads in 10µM levamisole in 1XM9, without bacteria. Oocytes from one gonad arm were imaged from each worm using a Nikon Eclipse TE2000-S inverted microscope equipped with a Plan Apo 60x/1.25 numerical aperture oil objective (Nikon Instruments Inc., Melville NY, USA). Images were captured using a Q imaging Exi Blue camera (Teledyne Photometrics, Tucson AZ, USA) using Nomarski optics and Q Capture Pro 7 software (Teledyne Photometrics, Tucson AZ, USA). The area of each fully cellularized oocyte within one gonad arm was measured in FIJI using the freehand tool with a scale of 4.64in by 3.47in (Schindelin et al. 2012). The area was measured by tracing around the membrane of each of the fully cellularized oocytes. Oocytes were considered fully cellularized if they had a cell membrane fully enclosing the nucleus. Images were taken in at least three biological replicates per strain at each temperature for a total of n between 56 and 179 cellularized oocytes across 14 to 18 worms per strain per temperature. Statistical analysis was done using a Two-Way ANOVA with Tukey’s multiple comparisons using Prism 10 (Graphpad, San Diego CA, USA).

### Lipid content analysis

Worms were maintained at 20°C until the L4 stage, at which point L4 worms were transferred to an NGM plate and placed at either 20°C, 26°C, or 27°C for 24 hours. P0 adults were transferred without bacteria to a 1XM9 solution on slides coated with poly-L-Lysine and embryos were extracted. Slides were fixed with 4% formaldehyde for 5 min then freeze-cracked in liquid Nitrogen. Slides were incubated in 1µg/mL Bodipy FL and 5mg/mL DAPI in 1XM9 for 1 hour at room temperature in the dark. Slides were washed three times with 1XM9 for 10 min in the dark and mounted on gelutol mounting medium. Images were captured using a Nikon A1R laser-scanning confocal unit controlled by NIS-Elements fitted on a Nikon inverted Eclipse Ti-E microscope with a Nikon DS-Qi1Mc camera and Plan Apo 60×/1.4 numerical aperture water objective. Images were taken in at least three biological replicates per strain at each temperature for a total of n between 17 and 31 embryos per strain per temperature. Images were scored using FIJI (Schindelin et al. 2012). Images were converted to 8-bit and the fluorescent threshold was set at 80-255. Individual droplets were separated via a watershed, and the number of individual lipid droplets were counted via analyzing particles restricted to the embryo. Normality of the data was assessed using a Shapiro-Wilk test and the data were found to be not normally distributed (p=8.499e-11). We used a Kruskal-Wallis test followed by a nonparametric pairwise Wilcoxon t-tests with the Bejamini-Hochberg correction to determine which data were significantly different from each other. All analysis was performed in RStudio (2024.12.1+563) using R-4.5.0 (R Core Team 2025).

### Length of L1s at hatching

Worms were maintained at 20°C until the L4 stage, at which point L4 worms were transferred to an NGM plate and placed at either 20°C, 26°C, or 27°C for 24 hours. After 24 hours, P0 adults were placed individually into 1XM9 solution without bacteria overnight at the appropriate temperature. Worms incubated at 26°C or 27°C for all strains were cut open to extrude embryos. F1 L1 larvae were transferred to slides coated with poly-L-Lysine. The entire body of L1 larvae were imaged from each worm using a Nikon Eclipse TE2000-S inverted microscope equipped with a Plan Apo 40x/1.4 numerical aperture air objective (Nikon Instruments Inc., Melville NY, USA). Images were captured using a Q imaging Exi Blue camera (Teledyne Photometrics, Tucson AZ, USA) using Nomarski optics and Q Capture Pro 7 software (Teledyne Photometrics, Tucson AZ, USA). The length of each L1 was measured using FIJI (Schindelin et al. 2012) using the freehand line tool to measure from the nose to the end of the tail with a scale of 3530 pixels/millimeter. Images were taken in at least three biological replicates per strain at each temperature for a total of n between 35 and 75 L1s across 6 to 17 worms per strain per temperature. Statistical analysis was done using a Two-Way ANOVA with Tukey’s multiple comparisons using Prism 10 (Graphpad, San Diego CA, USA).

## Data Availability Statement

Strains are available upon request. All raw data and images are available at XXXXX* *Full archive will be available for publication. All raw data was provided as supplemental spreadsheets names supplemental files Fig 1-6 and S1-S11 for review.

**Fig. 1:**
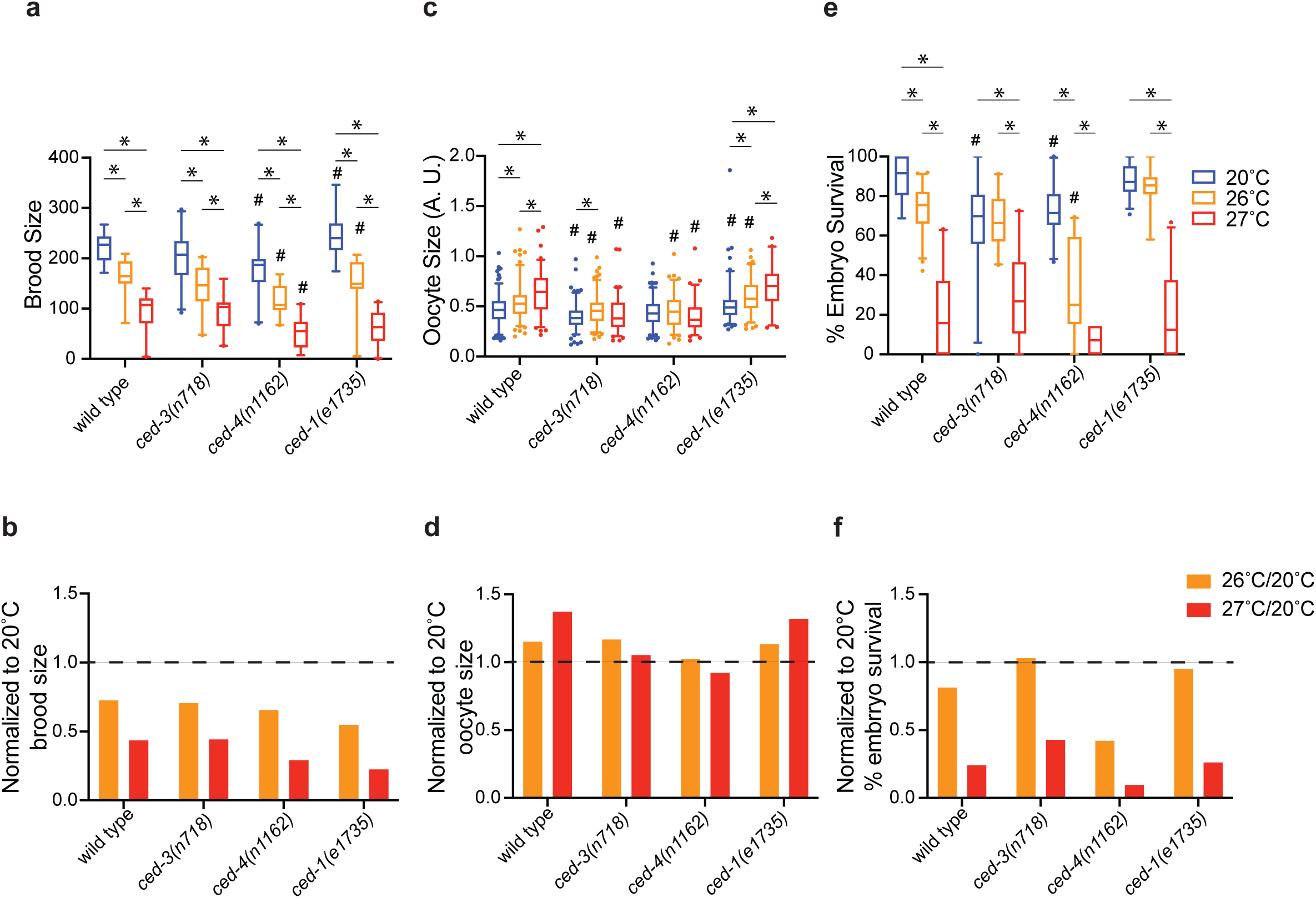
Fertility and embryo survival decrease and oocytes are smaller in mutants with no apoptosis. All experiments were done using wild type (N2), *ced-3(n718), ced-4(n1162),* and *ced-1(e1735)* mutants. For each experiment, hermaphrodites were either maintained continually at 20°C (blue) or upshifted to 26°C (orange) or 27°C (red) for 24 hours starting at the L4 stage. a) Fertility level was scored in n=18 to 21 worms per genotype per temperature. b) Fertility level normalized to 20°C at 26°C (orange) or 27°C (red). c) Size of cellularized oocytes in one gonad arm scored in n= 56 to 179 oocytes across 14 to 18 worms per genotype per temperature. d) Oocyte size normalized to 20°C at 26°C (orange) or 27°C (red). e) Percentage of hatched embryos scored in n=20 to 49 worms per genotype per temperature, with the exception of *ced-4(n1162)* at 27°C where only 4 worms were scored across 3 experiments because 81% of animals failed to lay any embryos. f) hatched embryos normalized to 20°C at 26°C (orange) or 27°C (red). Box plots represent 5-95% with center line at median and dots representing individual measurements outside the 5-95% range. Dashed line represents normalized level at 20°C. *p ≤ 0.05 significantly different within genotype between temperatures, #p ≤ 0.05 significantly different compared to wild type at same temperature using 2-way ANOVA with Tukey correction.

**Fig. 2:**
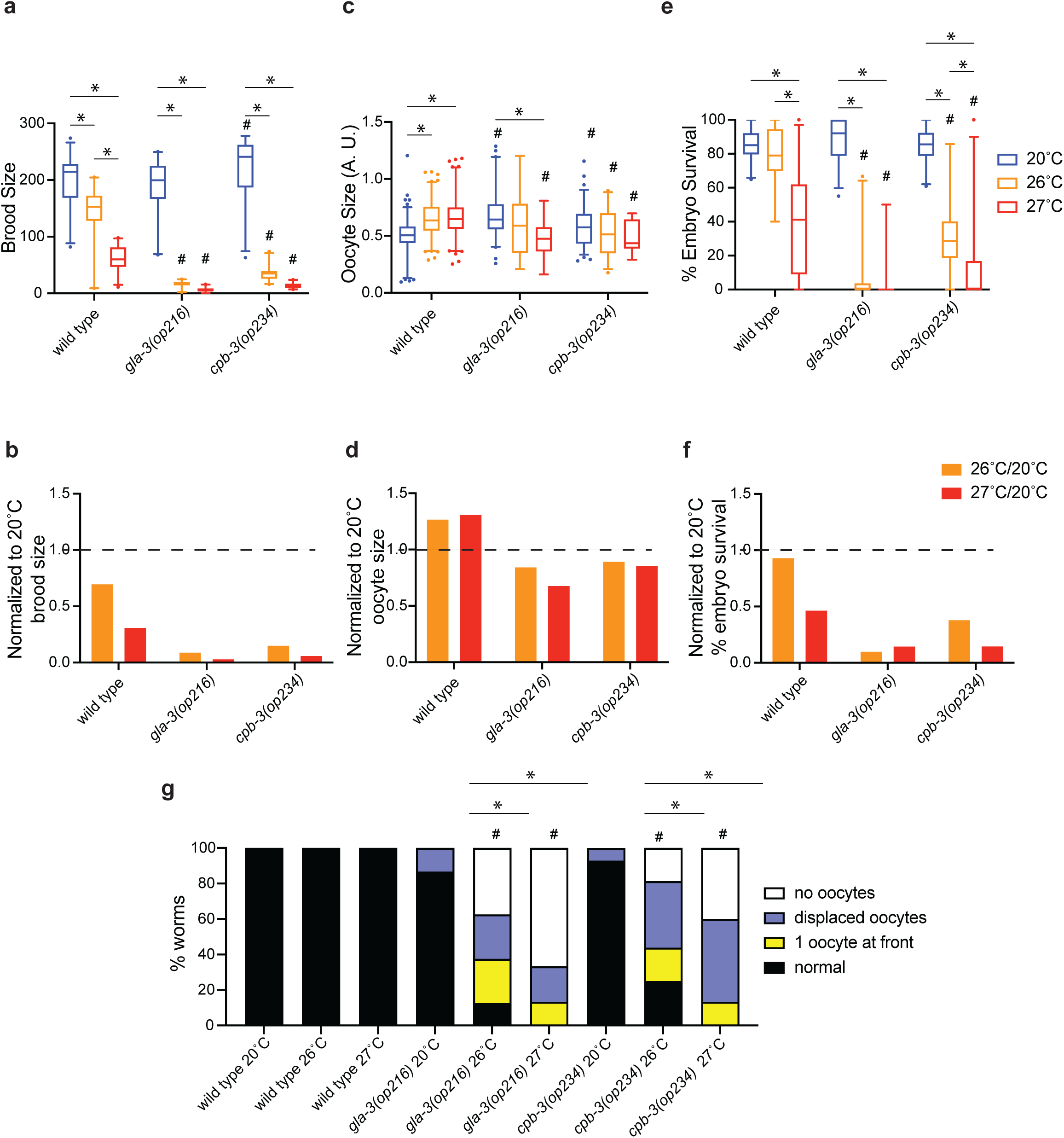
Fertility and embryo survival decrease and oocytes are smaller in mutants with elevated apoptosis during temperature stress. All experiments were done using wild type (N2), *gla-3(op216)*, and *cpb-3(op234)* mutants. For each experiment, hermaphrodites were either maintained continually at 20°C (blue) or upshifted to 26°C (orange) or 27°C (red) for 24 hours starting at the L4 stage. a) Fertility scored in n=20 to 24 worms per genotype per temperature. b) Fertility level normalized to 20°C at 26°C (orange) or 27°C (red). c) Size of cellularized oocytes in one gonad arm scored in n=11 to 116 oocytes across 14 to 16 worms per genotype per temperature. d) Oocyte size normalized to 20°C at 26°C (orange) or 27°C (red). e) Percentage of hatched embryos scored in n=9 to 26 worms per genotype per temperature. f) Percentage of hatched embryos normalized to 20°C at 26°C (orange) or 27°C (red). g) Percentage of worms with normal oocytes (black), 1 oocyte at front (yellow), displaced oocytes (orchid), or no oocytes (white) using the same worms as in (e). Box plots represent 5-95%, with center line at the median and dots representing individual measurements outside the 5-95% range. Dashed line represents normalized level at 20°C. *p≤0.05 significantly different within genotype between temperatures. #p ≤0.05 significantly different compared to wild type at same temperature using 2-way ANOVA with Tukey correction (a, c, e) or Fisher’s Exact test (g).

**Fig. 3:**
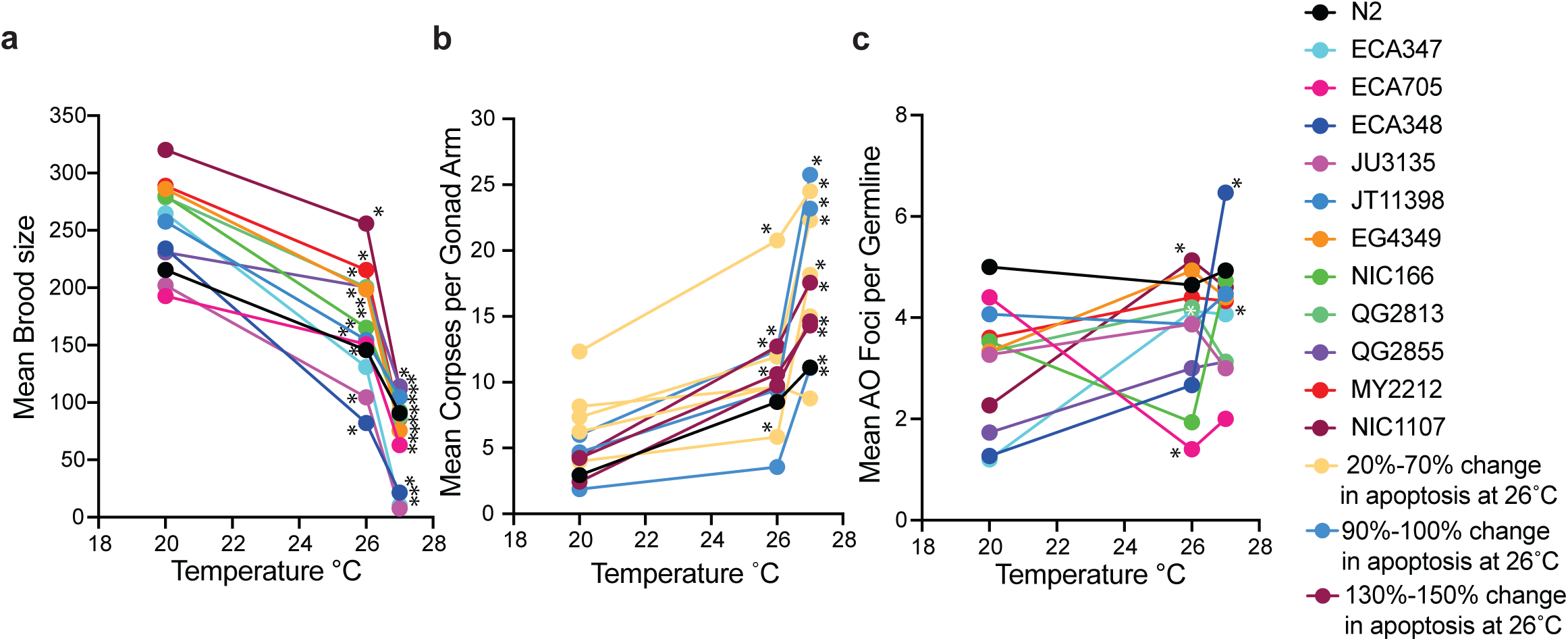
Fertility decreases and apoptosis increases in wild strains under a moderate temperature stress. All experiments were done using N2 and wild isolate strains listed in strain table. For each experiment, hermaphrodites were either maintained continually at 20°C or upshifted to 26°C or 27°C starting at the L4 stage. a) Mean brood size was scored in n=20 to 38 worms per strain per temperature. b) Apoptosis was scored using the *ced-1::gfp* transgene in n= 15 to 24 worms per strain per temperature. Strains were colored by their % change of apoptosis measured using the *ced-1::gfp* transgene at 26°C compared to 20°C. Small % change in apoptosis (gold) had an increase in apoptosis of 20% – 70%. Medium % change in apoptosis (aqua) had an increase in apoptosis of 90% - 100%. Large % change in apoptosis (magenta) had an increase in apoptosis of 150% - 300%. N2 (black) was measured in all assays. c) Apoptosis was scored using Acridine Orange dye in n=15 to 17 worms per strain per temperature. Each dot represents the mean measurement of a strain at each temperature. *p≤0.05 significantly different within strain between temperatures using 2-way ANOVA with Tukey correction.

**Fig. 4:**
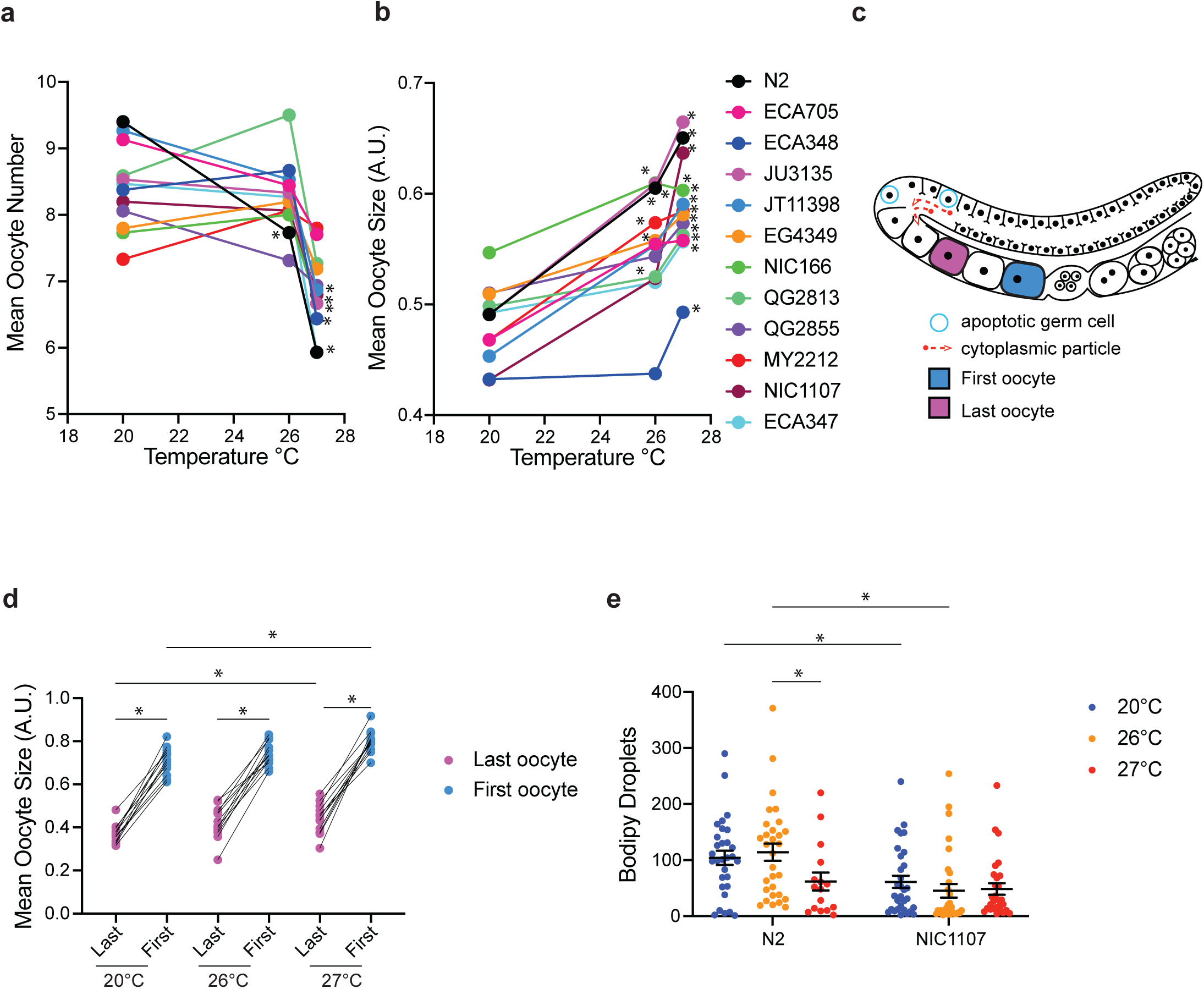
Oocytes are larger in wild strains under temperature stress. All experiments were done comparing N2 and wild strains listed in strain table. For each experiment, hermaphrodites were either maintained continually at 20°C or upshifted to 26°C or 27°C for 24 hours starting at the L4 stage. a) Number of cellularized oocytes were counted in one arm of the germline in n=14 to 18 worms per strain per temperature. b) Using the same worms as in (a), size of cellularized oocytes were measured in one arm of the germline in n=85 to 155 oocytes across 14 to 18 worms per strain per temperature. c) Schematic showing most recently cellularized oocyte (pink) and next oocyte to be ovulated (blue), designated as “first” and “last” oocytes with respect to the spermatheca. d) Size of last (pink) and first (blue) oocyte at 20°C, 26°C, and 27°C using same worms as in (b). e) number of lipid droplets in n=17 to 31 embryos per strain per temperature. Dots represent bodipy droplet in individual embryos with mean and error bars +/- SEM. *p≤0.05 significantly different within genotype between temperatures using 2-way ANOVA with Tukey correction (a and b) or Kruskal-Wallis test followed by Wilcoxon t-tests with Benjamini-Hochberg correction (e).

**Fig. 5:**
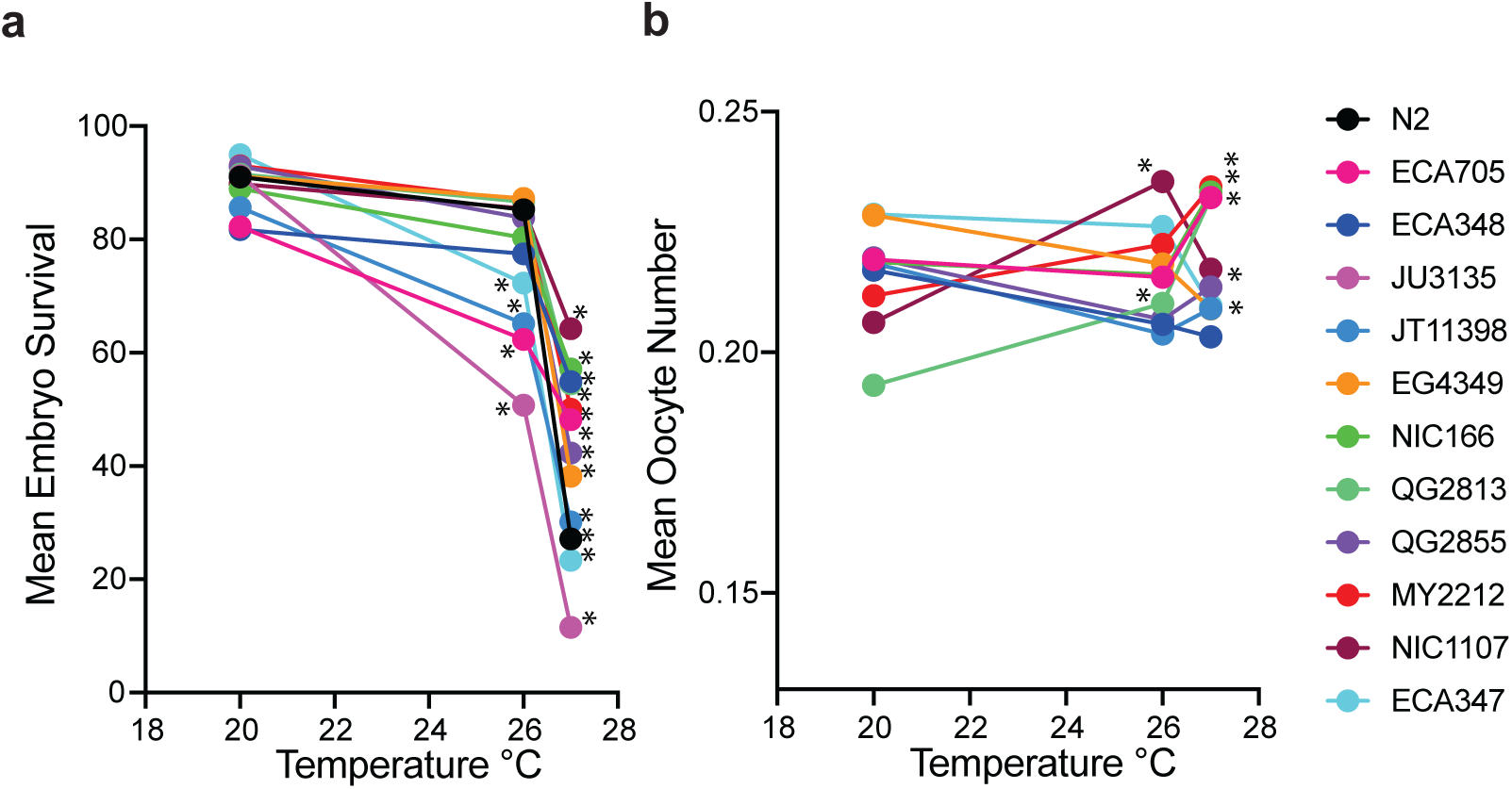
Embryo survival decreases and change in L1 length is inconsistent under temperature stress. All experiments were done using N2 and wild strains listed in strain table. For each experiment, hermaphrodites were either maintained continually at 20°C or upshifted to 26°C or 27°C for 24 hours starting at the L4 stage. a) Percentage of embryo survival scored in n=20 to 22 worms per strain per temperature. b) Length of L1 worms scored in n= 35 to 75 L1s which were progeny of 6 to 17 adults per strain per temperature. *p≤0.05 significantly different within genotype between temperatures using 2-way ANOVA with Tukey correction.

**Fig. 6:**
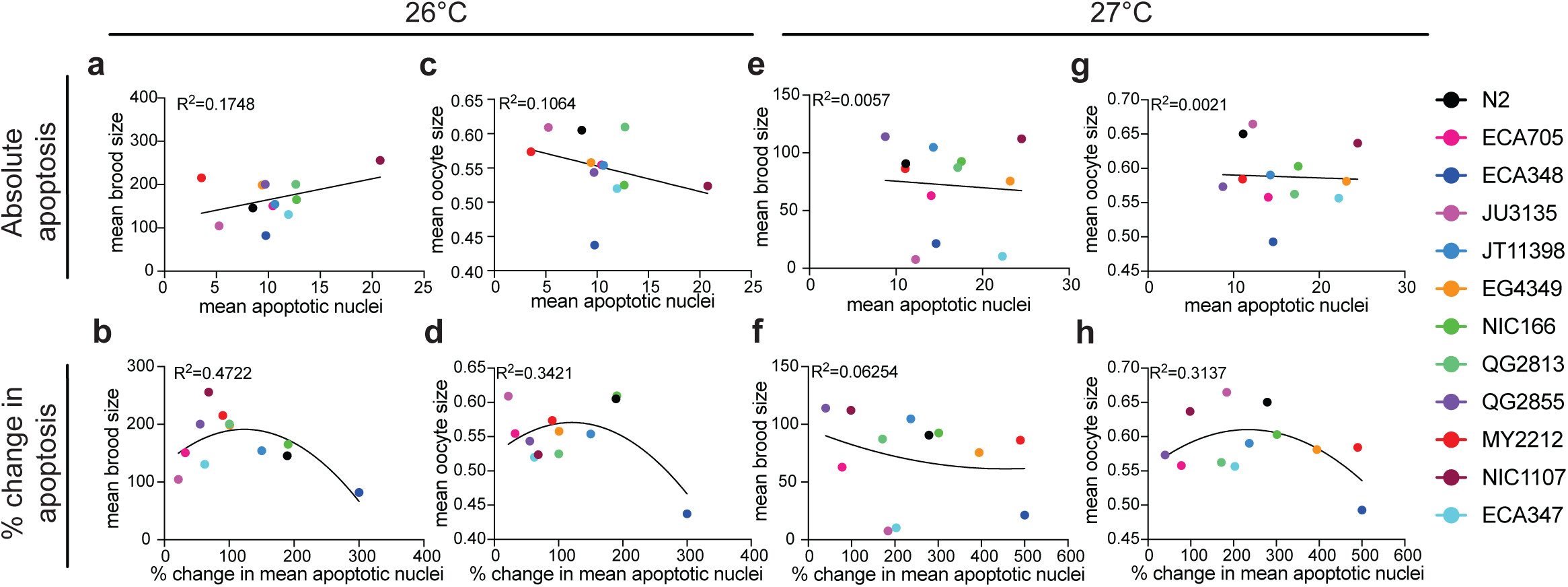
Intermediate % change in apoptosis is associated with highest fertility and oocyte size. Correlations were created using *ced-1::gfp* apoptosis data (Fig. 3b), brood size (Fig. 3) and oocyte size (Fig. 4) at 26°C and 27°C with linear regression (a, c, e, g) or second order polynomial regression (b, d, f, h). a) Mean brood size vs mean number of apoptotic nuclei at 26°C. P=0.1762. b) Mean brood size vs % change in mean number of apoptotic nuclei at 26°C. P=0.056. c) Mean oocyte size vs mean number of apoptotic nuclei at 26°C. P=0.3008. d) Mean oocyte size vs % change in mean number of apoptotic nuclei at 26°C. P=0.1520. e) Mean brood size vs mean number of apoptotic nuclei at 27°C. P=0.8159. f) Mean brood size vs % change in mean number of apoptotic nuclei at 27°C. P=0.7478. g) Mean oocyte size vs mean number of apoptotic nuclei at 27°C. P=0.8884. h) Mean oocyte size vs % change in mean number of apoptotic nuclei at 27°C. P=0.1838.

## Results

### Mutants with no apoptosis trended lower fertility, smaller oocytes, and decreased embryo survival under moderate temperature stress

To determine if apoptosis functions in the germline to maintain fertility and/or progeny fitness under temperature stress, we investigated how the loss of apoptosis or apoptotic cell engulfment affected fertility, oocytes, and embryo survival. We used *ced-3(n718)* and *ced-4(n1162)* mutants, which have no somatic or germline apoptosis (Yuan and Horvitz 1992; Shaham et al. 1999), and *ced-1(e1735)* mutants, which fail to engulf apoptotic nuclei into the surrounding somatic sheath cells (Zhou et al. 2001). All experiments were conducted at 20°C (non-stress), and at 26°C and 27°C (moderate temperature stress), since previous work has shown a significant induction in apoptosis (Compere et al. 2025) and decrease in fertility (Petrella 2014) at these temperatures. If apoptosis functions under moderate temperature stress to maintain fertility, then we would predict that *ced-3(n718)* and *ced-4(n1162)* mutants would have lower fertility compared to wild type. Because in *ced-1(e1735)* mutants apoptosis still occurs, but the corpses fail to be engulfed (Zhou et al. 2001), it is less clear what the effects of this mutation on fertility may be. We found that *ced-4(n1162)* mutants had significantly smaller brood size at 20°C, 26°C, and 27°C compared to wild type at the same temperature, while *ced-3(n718)* trended lower fertility but showed no significant difference in fertility compared to wild type at the same temperature (Fig. 1a and S1a). On the other hand, *ced-1(e1735)* had larger brood size at 20°C compared to wild type and smaller brood size at 26°C compared to wild type at the same temperature (Fig.1a and S1a). Consistent with previous studies, both wild type and mutant strains had a decrease in brood size at 26°C, and a larger decrease at 27°C compared to the same strain at 20°C (Cherian *et al*. 2020; Mikeworth *et al*. 2023; Fig. 1a). Because *ced-4(n1162)* showed differences in fertility even at 20°C, we normalized the brood size at 26°C and 27°C to the brood size at 20°C within the same genotype, thus allowing us to compare the strength of the effect of high temperature on fertility across strains. We found that *ced-4(n1162)* and *ced-1(e1735)* were more impacted by the effects of temperature than the other genotypes, especially at 27°C (Fig. 1b). Overall, this data suggests that the activation of apoptosis and engulfment are only modestly needed for maintaining a wild type level of progeny during temperature stress. is important for full brood size at all temperatures.

If apoptosis functions under moderate temperature stress to supply additional cytoplasmic resources to the remaining oocytes (Compere et al. 2025), we would predict that *ced-3(n718)* and *ced-4(n1162)* mutant oocytes would be smaller than wild type oocytes during temperature stress. We found that both *ced-3(n718)* and *ced-4(n1162)* mutants had smaller oocytes at 26°C and 27°C compared to wild type at the same temperature, while only *ced-3(n718)* had smaller oocytes at 20°C (Fig. 1c and S1b). On the other hand, *ced-1(e1735)* had larger oocytes at 20°C and 26°C compared to wild type at the same temperature (Fig. 1c and S1b). Consistent with previous studies, wild type, *ced-3(n718)*, and *ced-1(e1735)* mutants had larger oocytes at 26°C than at 20 °C, while only wild type and *ced-1(e1735)* had a larger oocytes at 27°C (Fig. 1c; Compere *et al*. 2025). Like with brood size, we normalized the size of oocytes at 26°C and 27°C to the size of oocytes at 20°C. For wild type and *ced-1(e1735)* mutants we found that oocytes had a continuing increase in size compared to 20°C as temperature increased, in *ced-3(n718)* mutants oocytes had an increase in size at 26°C only, while *ced-4(n1162)* show no increase in size at either temperature and even a small decrease in size at 27°C (Fig. 1d). We also counted the number of cellularized oocytes and found all mutants had significantly fewer oocytes than wild type at 20°C, *ced-4(n1162)* and *ced-1(e1735)* had significantly fewer oocytes at 26°C, and only *ced-1(e1735)* had significantly fewer oocytes at 27°C (Fig. S2). All genotypes were similarly impacted by temperature when the data were normalized to the number of oocytes at 20°C (Fig. S2). Overall, this data suggests that the activation of apoptosis is necessary for ensuring oocytes receive enough cytoplasmic resources at all temperatures. In addition, engulfment of apoptotic cells is generally dispensable for ensuring cytoplasmic resources are allocated into developing oocytes.

If apoptosis functions under a moderate temperature stress to maintain progeny fitness, we would predict that *ced-3(n718)* and *ced-4(n1162)* mutants would have lower embryo survival than wild type oocytes during temperature stress, while *ced-1(e1735)* mutants would have little to no effect on embryo survival. We found that both wild type and mutants showed a decrease in embryo survival at 26°C compared to 20°C, but the decrease was only significant in wild type and *ced-4(n1162)* mutants (Fig. 1e and S1c). On the other hand, all genotypes showed a significant decrease in embryo survival at 27°C compared to 20°C (Fig. 1e and S1c). When we normalized the 26°C and 27°C embryo survival to the 20°C embryo survival we found that were was a larger decrease in embryo survival only in *ced-4(n1162)* mutants at 26°C and 27°C (Fig. 1f). The embryo survival data, similar to the brood size and oocyte size data, shows that *ced-4(n1162)* mutants have a stronger effect on each metric at elevated temperatures than *ced-3(n718)* mutants. We also found that both *ced-3(n718)* and *ced-4(n1162)* mutants had significantly lower embryo survival at 20°C compared to wild type at 20°C (Fig. 1e). Since these mutants have no apoptosis, damaged nuclei that would normally be removed via apoptosis are likely being fertilized and laid as embryos but fail to hatch. The overall fertility level of these mutants is only slightly compromised while embryo survival is significantly decreased. Even though all genotypes laid similar number of embryos at 20°C (Fig. S2c), *ced-4(n1162)* had significantly lower brood size on day 2 compared to wild type (Fig. S2d), suggesting that the slight decline in fertility at 20°C in mutants with no apoptosis is driven by a decrease in embryo survival and not by an overall fewer number of progeny produced. Similarly at 26°C, the significant reduction in brood size in the *ced-4(n1162)* mutant is likely driven by a decrease in embryo survival as *ced-4(n1162)* mutant worms had significantly fewer progeny on day 1 and 2 compared to wild type (Fig. S2e) and a significant reduction in embryo survival (Fig. 1e).

However, the *ced-4(n1162)* mutant worms also laid fewer embryos at 26°C, suggesting that an overall lower production of embryos also contributes to the lower fertility observed (Fig. S2c). At 27°C, very few *ced-4(n1162)* worms laid any embryos, indicating that the significant decrease in brood size on day 1 and 2 in the *ced-4(n1162)* mutant is likely driven by an overall reduction in embryo production (Fig. S2c and S2f). Overall, this data suggests that the activation of apoptosis is necessary for maintaining a high levels of embryo survival at low temperatures. In addition, engulfment of apoptotic cells is generally dispensable for maintaining levels of embryo survival.

### Mutants with increased apoptosis had lower fertility, smaller oocytes, and decreased embryo survival under moderate temperature stress

Since the presence of apoptosis in the germline leads to slightly higher fertility, and better progeny fitness at all temperatures (Fig. 1), we investigated how increased apoptosis affected fertility, oocytes, and embryo survival. We used *gla-3(op216)* and *cpb-3(op234)* mutants, which have increased germline apoptosis under non-stress conditions, likely through two different mechanisms (Kritikou et al. 2006; Singh et al. 2017). If increased levels of apoptosis are protective during temperature stress, then *gla-3(op216)* and *cpb-3(op234)* mutants that increase apoptosis levels could show higher level of fertility and/or progeny fitness compared to wild type. *gla-3(op216)* and *cpb-3(op234)* mutants showed an equal or better brood size at 20°C compared to wild type (Fig. 2a and S1a); however, both mutants show a large and significant decrease in brood size at 26°C and 27°C, which was significantly smaller than the brood size of wild type at the same temperature (Fig. 2a and S1a). When normalized to fertility at 20°C, we found that there was a larger decrease in brood size in *gla-3(op216)* and *cpb-3(op234)* at both 26°C and 27°C than wild type (Fig. 2b). Overall, this data on mutants suggests that mutants with increased apoptosis have a stronger negative effect on fertility at elevated temperatures.

Given the effects of increased apoptosis on brood size during temperature stress, we wanted to know if there were similar detrimental effects on progeny fitness in high apoptosis mutants. We found that neither *gla-3(op216)* nor *cpb-3(op234)* mutants showed the increase in oocyte size with temperature that is seen in wild type (Fig. 2c and S1b) and in fact when oocyte size was normalized to 20°C the trend showed that *gla-3(op216)* and *cpb-3(op234)* had smaller oocytes during temperature stress (Fig. 2d). The decrease in oocyte size during temperature stress in *gla-3(op216)* and *cpb-3(op234)* mutants was accompanied by a dramatic decrease in oocyte number at 26°C and 27°C (Fig. S3a). We also found that *gla-3(op216)* and *cpb-3(op234)* mutants had lower embryo survival at both 26°C and 27°C compared to wild type at the same temperature (Fig. 2e and S1c) and when embryo survival is normalized to 20°C both mutants showed a much stronger effect of temperature on embryo survival than wild type (Fig. 2f). The low embryo survival seen in g*la-3(op216)* mutants was exacerbated by the low number of embryos that the strain laid at elevated temperatures (Fig. S3c). In addition, we observed a significantly higher frequency of disorganized germline patterning in *gla-3(op216)* and *cpb-3(op234)* specifically under temperature stress (Fig. 2g). When we used 18 hours of temperature stress instead of 24 hours, we found no significant changes in oocyte size when comparing 18 hour and 24-hour stress (Fig. S4). Overall, this data suggests that high levels of apoptosis leads to a disorganized germline pattern, which may contribute to the decrease in fertility and progeny fitness we observed.

### Apoptosis increases under a moderate temperature stress in wild strains of C. elegans

Previous work has shown there is a range in the level of disruption of fertility in recently isolated strains of *C. elegans* when exposed to a moderate temperature stress (Petrella 2014; Zhang et al. 2021). We wanted to investigate if there was a similar range in the level of apoptosis increased during moderate temperature stress across wild strains of *C. elegans.* We selected 11 wild strains isolated from the environment that show a range in fertility level under temperature stress (Fig. 3a, S1a, and S5). Using those strains and the N2 wild type, we created near isogenic lines with the *ced-1::gfp* transgene crossed into each of the wild strains and our lab’s N2. Similar to what has been previously found in the N2 background (Poullet et al. 2015; Compere et al. 2025), we found that five strains had a significant increase in apoptosis at 26°C, and 10 strains had a significant increase in apoptosis at 27°C (Fig. 3b and S6). To more easily visualize the induction of apoptosis, we colored each wild strain according to its % change in apoptosis at 26°C. Five strains were classified as a small % change (20%-70% increase), three strains were classified as a medium % change (90%-100% increase), and three strains were classified as a large % change (150%-300%). To investigate the relationship between the level of apoptosis and the induction of apoptosis during temperature stress, we looked for a correlation between the number of apoptotic nuclei and either the absolute increase in apoptotic nuclei or the % change in apoptotic nuclei between 20°C and either 26°C or 27°C (Fig. S7). We found a significant positive correlation at both 26°C and 27°C between the mean apoptotic nuclei in each wild strain and the absolute increase in apoptosis at the respective temperature, suggesting that strains with more apoptosis during temperature stress have a larger absolute increase in apoptosis (Fig. S7). While not a significant relationship, the trendlines at both 26°C and 27°C when comparing the number of apoptotic nuclei with the % change in apoptosis suggest that strains with an intermediate amount of apoptosis have the largest % change in apoptosis (Fig. S7).

We next wanted to measure apoptosis without the use of a transgene. So, we used Acridine Orange dye as an alternative method. We found that only two wild strains had significantly increased apoptosis under a moderate temperature stress using this method for assessing apoptosis (Fig. 3c and S6). Instead, most strains, including N2, showed no significant change in apoptosis, and one strain had significantly decreased apoptosis under a moderate temperature stress compared to non-stress conditions (Fig. 3c and S6). With both methods of evaluating apoptosis there was a range of apoptotic cells between the strains with some strains having more apoptotic cells compared to others at the same temperature (Fig. 3b, c, and S6).

When apoptosis is normalized to the physiological level, we observe a range in the effect of temperature in both methods of apoptosis (Fig. S6). As a third method we also used *act-5::yfp* transgene in an N2 background to measure apoptosis under temperature stress and found a significant increase in apoptosis at 27°C (Fig. S8). Overall, these data suggests that apoptosis, when measured by AO, does not increase in most wild strains tested and N2, but does show an increase when measured using the *ced-1::gfp* or *act-5::yfp* transgene.

### Cellularized oocytes increase in size under moderate temperature stress

To determine if there was a correlation between the level of apoptosis and partitioning of resources in oocytes in wild strains and N2, we measured oocytes in hermaphrodites that were upshifted to 26°C and 27°C. When analyzing if there was a decrease in oocyte numbers under temperature stress, we found that at 26°C only N2 has a significant decrease in the number of cellularized oocytes, while at 27°C half of the strains showed a significant decrease in the number of cellularized oocytes (Fig. 4a and S9). The other strains showed a non-significant trend of decreased number of oocytes under a temperature stress (Fig. 4a and S9). When analyzing oocyte size, we found that most strains had larger oocytes at 26°C, and all strains had larger oocytes at 27°C (Fig. 4b and S9). When normalized to oocyte size at 20°C, we found that strains had larger oocytes at 26°C, and some strains further increased in size at 27°C, while others showed no additional size increase (Fig. S9). These data suggest that for most strains there were larger oocytes, and sometimes fewer oocytes, under a moderate temperature stress, and that temperature has a greater effect on the size of oocytes in some wild strains and N2.

Previous studies have also shown that the increase in size of oocytes may be due to increased cytoplasmic streaming under a moderate temperature stress rather than growth after the cellularization process (Wolke et al. 2007; Compere et al. 2025). We compared the size of the most recently cellularized oocyte and the size of the oocyte next to be ovulated using the data from Fig. 4a (Fig. 4d and S9). We found that there was a significant increase in the size of the next oocyte to be ovulated compared to the most recently cellularized oocyte within each temperature, suggesting a role of a post-cellularization growth mechanism in the increased size of oocytes during elevated temperatures. We also found that both the most recently cellularized oocyte and the next oocyte to be ovulated was significantly larger at 27°C compared to 20°C (Fig. 4d and S9), suggesting that increased streaming contributes to the size growth at 27°C. These data all together suggest that both cytoplasmic streaming and post-cellularization growth significantly contribute to the overall increase in size in oocytes under temperature stress across N2 and newly isolated wild strains.

To investigate a possible method of post-cellularization growth we measured the number of lipid droplets using Bodipy in early embryos in N2 and the newly isolated wild strain NIC1107 to evaluate changes in importation of yolk lipoproteins from the intestines. NIC1107 was chosen as an example of a newly isolate strain because if its large number of apoptotic cells (Fig. 3b), and larger brood size (Fig. 3a) under a moderate temperature stress. We found no significant change in the number of lipid droplets as temperature increased in either strain (Fig. 4e). There were significantly more lipid droplets at 20°C and 26°C in N2 compared to NIC1107 (Fig. 4e). The lack of increased bodipy staining indicates that the increase in size of cellularized oocyte may not be due to an increase in yolk lipoproteins added post-cellularization and may be due to another method of post-cellularization growth. Overall, these results suggest that oocytes in newly isolated wild strains and N2 are impacted by the effects of temperature stress by increasing in size, potentially to supply early embryos with additional cytoplasmic nutrients.

### Embryo survival decreases and Length of L1s do not change consistently under moderate temperature stress

Since we have shown an effect of temperature on developing oocytes inside the hermaphrodite, we wanted to determine the effects on the progeny of hermaphrodites that experience temperature stress. First, we evaluated embryo survival under a moderate temperature stress. We found that three strains had a significant decrease in embryo survival at 26°C and all strains has a significant decrease at 27°C (Fig. 5a and S10). Of the strains with non-significant changes at 26°C, all showed a trend of lower embryo survival (Fig. 5a and S10). When normalized to the rate of embryo survival at 20°C, we found that strains that also have lower fertility showed a larger decrease in hatched embryos (Fig. S5 and S10). Strains that had smaller brood sizes during moderate temperature stress were also strains to show lower embryonic survival, especially at 26°C (Fig. S5 and S10). These data suggest that embryo survival decreases under a moderate temperature stress in wild strains and likely contributes to the lower fertility observed in some strains.

As a second measure of progeny fitness, we measured the length of L1 worms laid from hermaphrodites that were exposed to a moderate temperature stress. At 26°C, we found that only QG2813 and NIC1107 showed an increase in L1 length (Fig. 5b and S10). While at 27°C, we found that two different strains, ECA347 and EG4349, had a significant decrease in L1 length, and three strains, NIC166, QG2813, and MY2212 had a significant increase in the length of L1s compared to 20°C (Fig. 5b and S10). When normalized to length at 20°C, we found there was no consistent effect of temperature on length of L1 across strains (Fig. S10). These data suggest that moderate temperature stress may not have a consistent impact on the length of L1 worms in wild strains, and that the size of L1s may be limited by other factors, such as the size of eggshell. Overall, these data from all progeny fitness traits show that a moderate temperature stress does, in general, impact the progeny of mothers that experience temperature stress.

### Higher apoptosis weakly correlates with better fertility but smaller oocytes under moderate temperature stress

To more directly understand if there is a relationship between apoptosis and the life history traits we measured during moderate temperature stress (brood size, oocyte size, embryo survival, and L1 length), we looked for correlations between either mean apoptosis levels or the % change in apoptosis with temperature stress and these other measures. We found no relationship between the number of apoptotic nuclei and the life history traits at any temperature (Fig. 6 and S11). However, at 26°C for comparisons with % change in apoptosis we found no statistically significant relationships, we did identify an overall trend that strains with an intermediate % change in apoptosis had the highest fertility, oocyte size, and embryo survival at 26°C (Fig. 6 and S11). On the other hand, at 27°C we found overall trends that strains with an intermediate % change in apoptosis had the largest oocytes but the lowest embryo survival, indicating the severity of the effects of temperature stress just below sterility (Fig. 6h and S11k). The length of L1 worms showed no relationship with apoptosis, further indicating that the size of L1 worms may be limited by factors like eggshell size (Fig. 6 and S11). These correlation plots suggest that a small or large % increase in apoptosis in response to temperature stress may be less protective for fertility and progeny fitness than a more intermediate increase in apoptosis in response to temperature stress.

## Discussion

Oogenic nuclei have the need to both supply a full complement of cytoplasmic resources and an accurate genome to the developing embryos in both non-stress and stress conditions to produce the highest number of fit progeny. Here, we’ve shown that the presence of apoptosis at an intermediate level is necessary for the highest fertility level and most fit progeny in both unstressed and stressed conditions. Additionally, within a set of wild strains isolated from the environment, we demonstrate a range in both the amount and induction in apoptosis, brood size, oocyte features, embryo survival, and length of L1 worms, and that an intermediate induction of apoptosis leads to the highest fertility level and most fit progeny. Our findings support a model that apoptosis increases in the germline in response to temperature stress to buffer fertility and ensure progeny fitness. Our work expands on the knowledge on the response to a moderate temperature stress and enhances our knowledge of the role of apoptosis in promoting fertility and high-quality oocytes and progeny in both non-stress and stress conditions.

### Both the proper level of apoptosis and induction of apoptosis play a role in buffering the effects of moderate temperature stress

Our data suggests a model that starting with an intermediate absolute level of apoptosis in the germline is most beneficial for fertility, oocyte provisioning, and embryo survival during moderate temperature stress. A lack of apoptosis led to a more severe impact of temperature on fertility, oocytes, and embryo survival, indicating that the presence of apoptosis during stress is critical. Similar impacts on fertility and progeny fitness have been reported during acidic, oxidative, starvation, or alcoholic conditions in the absence of apoptosis (Salinas et al. 2006; Fausett et al. 2021), suggesting a role for the presence of apoptosis as a stress response mechanism in the germline to a variety of stressors. Our work is the first to show that under stress conditions mutants with high levels of apoptosis have significantly diminished fertility and progeny fitness (Fig. 2 and S1). It is likely that any increase in apoptosis on top of the already high basal level in these mutants depletes the germ cell pool required to supply cytoplasmic resources to developing oocytes. Our group of wild strains during temperature stress were generally in the middle in the effect of temperature on fertility, oocyte size, and embryo survival, further indicating an intermediate level of apoptosis is best during a moderate temperature stress. Together, these results support a model in which germline apoptosis operates within an optimal range in both the unstressed and the stressed germline: insufficient apoptosis fails to remove damaged nuclei and excessive apoptosis reduces cytoplasmic resources necessary for fit progeny.

Although *ced-3(n718)* and *ced-4(n1162)* mutants have both been shown to have no apoptosis (Salinas et al. 2006), *ced-4(n1162)* mutants consistently exhibited more severe defects than *ced-3(n718)* mutants under moderate temperature stress. Similar differences using *ced-3* and *ced-4* mutants have been reported when measuring aging and other stressors (Aballay and Ausubel 2001), suggesting distinct roles for CED-3 and CED-4 in relation to the downstream effects of apoptosis on fertility and progeny fitness. Additionally, *ced-3* mutants have been shown to confer a stronger resistance to stress in the soma compared to *ced-4* mutants (Judy et al. 2013), suggesting *ced-4* mutants may be more susceptible to the effects of temperature stress, contributing to the more severe defects we observed. Nonetheless, loss of either factor compromises fertility, oocyte partitioning, and embryo survival, especially in stress environments. Together, these finding reinforce the importance of the canonical apoptotic machinery in sustaining germline function, while also raising the possibility of distinct roles for individual pathway components.

In addition to the starting level of apoptosis being important for the buffering of moderate temperature stress, our data support a model that an induction of apoptosis at intermediate levels supplies additional cytoplasmic resources to the developing oocytes that ensures progeny fitness during stress. Numerous studies have demonstrated that germline apoptosis is induced above basal levels in response to environmental stressors, including temperature, starvation, and oxidative stress (Salinas *et al*. 2006; Poullet *et al*. 2015; Fernández-Cárdenas *et al*. 2017; Compere *et al*. 2025: this study). We’ve shown here that mutants with excess apoptosis above the physiological level during no stress have oocyte numbers and sizes comparable to temperature-stressed wild type animals (Fig. 2 and S1), suggesting that the induction of apoptosis above physiological levels supplies additional cytoplasmic resources to the developing oocytes. Previous work examining DREAM complex members LIN-35 and LIN-54 have shown these proteins are necessary for the full induction of apoptosis during temperature stress (Compere et al. 2025). *lin-35* and *lin-54* mutants have also been shown to have reduced fertility (Cherian et al. 2020; Mikeworth et al. 2023) and smaller cellularized oocytes (Compere et al. 2025) compared to wild type during temperature stress. Additionally, wild strains in our study with a medium % change in apoptosis had the highest level of fertility, largest oocytes, and highest embryo survival, while a small % change or large % change had lower fertility, smaller oocytes, and lower embryo survival (Fig. 6 and S11). An intermediate induction of apoptosis during stress balances the need to supply additional cytoplasmic resources to developing oocytes while keeping enough oogenic nuclei in the germline to produce those cytoplasmic resources. Taken together, these data suggest that insufficient induction of apoptosis compromises fertility, oocyte provisioning, and progeny fitness. At the same time, our analyses also indicate that a large induction of apoptosis may similarly impact fertility, oocytes, and progeny fitness, likely through depletion of germ cells in the germline as seen in mutants with excess apoptosis that show severely diminished fertility, few or no oocytes, and the proximal expansion of pachytene-staged nuclei where cellularized oocytes normally occupy. Collectively, these observations support our model that an intermediate induction of apoptosis maximizes fertility and progeny fitness during temperature stress by removing damaged nuclei while maintaining enough germ cells to produce the additional cytoplasmic resources to ensure progeny are fit during environmental stress.

Our measurements of germline apoptosis using the *ced-1::gfp* reporter in the N2 wild type are consistent with previous reports of both physiological (Poullet et al. 2015; Fernández-Cárdenas et al. 2017; Compere et al. 2025) and temperature-stress induction of apoptosis (Poullet et al. 2015; Compere et al. 2025). However, we did not observe a comparable increase when apoptosis was assessed using acridine orange staining (Fig. 3). This discrepancy highlights an important methodological consideration: different assays capture distinct stages of the apoptotic process and may yield slightly different outcomes. The *ced-1::gfp* transgene labels apoptotic germ cells that have been recognized for engulfment by the somatic sheath cells and are in the early stages of apoptosis (Lant and Derry 2014). In contrast, acridine orange is considered a late stage marker of apoptosis as it stains condensed DNA and requires membrane permeability and nuclear condensation (Lant and Derry 2014). Since biological processes often increase in speed as temperature increases (Knapp and Huang 2022), an increase in apoptotic initiation combined with a more rapid corpse clearance could produce a detectable increase in CED-1::GFP-positive corpses without a proportional accumulation of acridine orange-positive nuclei. Importantly, previous work using *ced-1::gfp* transgene have demonstrated that increased signal under stress conditions reflect elevated apoptotic cell death rather than delayed engulfment (Salinas et al. 2006). While we cannot fully exclude subtle differences in these assays as reason for differences between *ced-1::gfp* and acridine orange, the consistency of *ced-1::gfp* transgene-based measurements of apoptosis supports the interpretation that temperature stress induces a genuine increase in germline apoptosis.

### Loss of regulation of apoptosis has different effects in the soma and germline during stress

There is only a small number of somatic cells that die post-embryonically through apoptosis and all of those deaths occur prior to the L4 stage in hermaphrodites (Sulston and Horvitz 1977). Therefore, the germline is the only location in adult hermaphrodites where apoptosis occurs (Gartner et al. 2008). Maybe unsurprisingly, there are opposite effects of stress on somatic and germline phenotypes when apoptosis is misregulated in apoptosis or engulfment defective mutants (Salinas et al. 2006; Schertel and Conradt 2007; Haskins et al. 2008; Judy et al. 2013; Poullet et al. 2015; Fausett et al. 2021; Wan et al. 2021; Compere et al. 2025; Salinas et al. 2026). Across numerous studies, mutants that disrupt the ability to carry out apoptosis, like in *ced-3* and *ced-4* mutants, induce apoptosis, like *lin-35* and *lin-54* mutants, or limit apoptosis, like *gla-3* mutants, lead to a decrease in stress buffering when looking at phenotypes associated with the germline, such as fertility or progeny fitness metrics. This higher level of stress sensitivity has been shown for many types of stress, including heat shock, osmotic stress, UV treatment, oxidative stress, acid stress, ethanol stress, starvation, and moderate temperature stress (Salinas et al. 2006; Schertel and Conradt 2007; Láscarez-Lagunas et al. 2014; Poullet et al. 2015; Fausett et al. 2021; Compere et al. 2025; Salinas et al. 2026; this study). However, mutants that disrupt either the ability to do apoptosis or engulf apoptotic cells, like *ced-1* mutants, show an increased ability to buffer stress when looking at phenotypes associated with the soma, in particular post-developmental survival (Judy et al. 2013; Wan et al. 2021). Interestingly, this decreased stress sensitivity in apoptosis and engulfment mutants was only associated with some stressors, such as osmotic stress, heat shock, ER stress, and some pathogens (Judy et al. 2013; Wan et al. 2021), but not UV treatment or oxidative stress (Judy et al. 2013). Our study complements previous work showing increased stress sensitivity in the germline during moderate temperature stress in apoptosis and engulfment deficient mutants, and adds that mutants with elevated apoptosis during moderate temperature stress also have increased stress sensitivity in the germline. Further work is needed to see if there is a link between the increase in stress sensitivity in the germline and decrease in stress sensitivity in the soma in apoptosis and engulfment deficient mutants as loss of apoptosis in the germline could free some cellular energy stores to be used to respond more strongly to stresses in somatic cells. Additionally, it would be interesting to investigate the effect in the soma of elevated apoptosis in the germline as the increased energy investment in the germline to carry out apoptosis could confer a weaker stress response in the soma.

### Phenotypic variation in wild strains may be ecologically relevant

In this study, we’ve characterized apoptosis, fertility, and multiple progeny fitness metrics across wild strains, including oocyte size, embryo survival and larval size in both an unstressed and an ecologically relevant stress condition: a moderate temperature stress. We’ve shown that each of these metrics have variation across the strains assayed, and N2 typically in the middle when compared across all strains (Fig. S1). This work expands upon other studies using wild strains have characterize the variation in a variety of phenotypes, like pathogen avoidance (Martin et al. 2017), starvation response (Webster et al. 2022), and temperature stress (Harvey and Viney 2007; Petrella 2014; Zhang et al. 2021) and the more to understand phenotypes across a broader range of genetics background outside of the canonical N2 laboratory reference strain. Understanding the phenotypic variation is important for understanding real world ecological consequences of stress in natural systems. Under stress conditions especially, populations of strains with traits like higher fertility, larger oocytes, and higher embryo survival could persist more easily than populations of strains with lower fertility and progeny fitness. Furthermore, strains with an intermediate amount and induction of apoptosis during temperature stress are more likely to persist during stress, due to the removal of damaged nuclei and supply of additional cytoplasmic resources.

Consistent with this idea, several strains examined in this study, NIC1107, QG2813, and MY2212, consistently maintained higher fertility, larger oocytes, more viable embryos, and intermediate apoptosis under temperature stress. These traits may allow these strains to better sustain reproduction during temperature stress, potentially providing a selective advantage. In contrast, strains such as JU3135, ECA348, and ECA347 exhibited reduced fertility, smaller oocytes, lower embryo survival, and low apoptosis, suggesting a greater sensitivity to temperature stress. Ultimately, populations that are more resistant to the detrimental effects of temperature stress are more likely survive and continue to produce offspring until more favorable environmental conditions arise.

Together, our findings highlight the importance of studying natural variation to understand how biological processes like germline apoptosis may be a strategy *C. elegans* uses to maintain the ability to produce fit progeny despite experiencing environmental stressors. By moving beyond the N2 reference strain and incorporating diverse wild strains, we can begin to uncover how evolutionary pressures shape stress responses in natural populations of *C. elegans*.

## Strain table

| Strain | Genotype | Source |
| --- | --- | --- |
| CB3203 | *ced-1(e1735) l* | CGC |
| ECA347 | Wild isolate | CaeNDR |
| ECA348 | Wild isolate | CaeNDR |
| ECA705 | Wild isolate | CaeNDR |
| EG4349 | Wild isolate | CaeNDR |
| JT11398 | Wild isolate | CaeNDR |
| JU3135 | Wild isolate | CaeNDR |
| LNP0002 | *petlR2(V;bcls39 [lim-7p::ced-1::GFP + lin-15(+)]*<br>*N2>JT11398)* | This study |
| LNP0003 | *petlR3(V;bcls39[lim-7p::ced-1::GFP + lin-15(+)]*<br>*N2>N2)* | (Zampetti et al. 2025) |
| LNP0006 | *petlR6(V;bcls39[lim-7p::ced-1::GFP + lin-15(+)]*<br>*N2>ECA348)* | This study |
| LNP0033 | *petlR14(V;bcls39[lim-7p::ced-1::GFP+lin-15(+)]*<br>*N2>MY2212)* | This study |
| LNP0036 | *petlR17(V;bcls39[lim-7p::ced-1::GFP+lin-15(+)]*<br>*N2>QG2855)* | This study |
| LNP0047 | *petlR20(V;bcls39[lim-7p::ced-1::GFP+lin-15(+)]*<br>*N2>NIC166)* | This study |
| LNP0052 | *PetlR8(V;bcls39[lim-7p::ced-1::GFP+lin-15(+)]*<br>*N2>ECA347)* | This study |
| LNP0055 | *petlR11(V;bcls39[lim-7p::CED-1::GFP+lin-15(+)]*<br>*N2>NIC1107)* | This study |
| LNP0069 | *petIR25(V;bcls39[lim-7p::CED-1::GFP+lin-15(+)])*<br>*N2>EG4349* | This study |
| LNP0118 | *petIR26(V;bcls39[lim-7p::CED-1::GFP+lin-15(+)])*<br>*N2>ECA705* | This study |
| LNP0121 | *petIR29(V;bcls39[lim-7p::CED-1::GFP+lin-15(+)])*<br>*N2>JU3135* | This study |
| LNP0124 | *petIR32(V;bcls39[lim-7p::CED-1::GFP+lin-15(+)])*<br>*N2>QG2813* | This study |
| MD701 | *bcls39 [lim-7p::ced-1::GFP + lin-15(+)]* | (Zhou et al. 2001;<br>Schumacher et al. 2005),<br>CGC |
| MT1743 | *ced-3(n718) IV* | CGC |
| MT2547 | *ced-4(n1162) III* | CGC |
| MY2212 | Wild isolate | CaeNDR |
| N2 | Wild type | Strome Lab |
| NIC1107 | Wild isolate | CaeNDR |
| NIC166 | Wild isolate | CaeNDR |
| QG2813 | Wild isolate | CaeNDR |
| QG2855 | Wild isolate | CaeNDR |
| WS1137 | *cpb-3(op234) I* | CGC |
| WS2170 | *opls110 [lim-7p::YFP::actin + unc-119(+)]* | CGC |
| WS2972 | *gla-3(op216) I* | CGC |

---

**Fig S1: Fertility, size of oocytes, and embryo survival decreased across all genotypes** Hermaphrodites were either maintained continually at 20°C or upshifted to 26°C or 27°C for 24 hours starting at the L4 stage. Data was combined from no-apoptosis mutants (lavender), *ced-1(e1735)* (green), high apoptosis mutants (salmon), wild strains (gray), and N2 wild type (black) for a) fertility (Fig. 1A, 2A, and 3A), b) oocyte size (Fig. 1C, 2C, 4B), and c) embryo survival (Fig. 1E, 2E, 5A). #p ≤0.05 mutants significantly different compared to wild type from same experiment at same temperature using 2-way ANOVA with Tukey correction.

**Fig. S2: Number of oocytes decreases, fewer embryos are laid, and brood size is slightly decreased in no-apoptosis mutants under moderate temperature stress** Hermaphrodites were either maintained continually at 20°C (blue) or upshifted to 26°C (orange) or 27°C (red) for 24 hours starting at the L4 stage. a) Number of oocytes were scored in wild type (N2), *ced-3(n718), ced-4(n1162),* and *ced-1(e1735)* mutants in n= 14 to 18 worms per genotype per temperature. b) Number of oocytes normalized to 20°C in wild type (N2), *ced-3(n718), ced-4(n1162),* and *ced-1(e1735)* mutants at 26°C (orange) or 27°C (red). Dashed line represents normalized level at 20°C. c) Embryos laid in six hours in wild type (N2), *ced-3(n718), ced-4(n1162),* and *ced-1(e1735)* mutants in n=20 to 49 worms per genotype per temperature, same worms as in Fig. 1e. Data from Fig. 1a separated by day in hermaphrodites maintained either continually at 20°C (a) or upshifted to 26°C (b) or 27°C (c) starting at the L4 stage in wild type (N2; blue), *ced-3(n718)* (green), *ced-4(n1162)* (red), and *ced-1(e1735)* (black) mutants. Fertility was scored in n=18 to 21 worms per genotype per temperature. Each point represents mean brood size with error bars +/- SEM. Box plots represent 5-95% with center line at median. *p≤0.05 significantly different within genotype between temperatures, #p ≤0.05 significantly different compared to wild type at same temperature using 2-way ANOVA with Tukey correction.

**Fig. S3: Number of oocytes decreases and fewer embryos laid in high-apoptosis mutants under moderate temperature stress.** a) Number of oocytes were scored in wild type (N2), *gla-3(op216)*, and *cpb-3(op234)* mutants. Hermaphrodites were either maintained continually at 20°C (blue) or upshifted to 26°C (orange) or 27°C (red) for 24 hours starting at the L4 stage. n= 14 to 16 worms per genotype per temperature. Box plots represent 5-95%, with center line at the median. c) Embryos laid in six hours in wild type (N2), *gla-3(op216)*, and *cpb-3(op234)* mutants in n=18-28 worms per genotype per temperature, same worms as in Fig. 2e. *p≤0.05 significantly different within genotype between temperatures. #p ≤0.05 significantly different compared to wild type at same temperature using 2-way ANOVA with Tukey correction. b) number of oocytes normalized to 20°C in wild type (N2), *gla-3(op216)*, and *cpb-3(op234)* mutants at 26°C (orange) or 27°C (red). Dashed line represents normalized level at 20°C.

**Fig. S4: 18hrs and 24hrs of stress have same effect on oocytes in high-apoptosis mutants** Hermaphrodites were either maintained continually at 20°C (blue) or upshifted to 26°C (orange) or 27°C (red) for 18 or 24 hours starting at the L4 stage. Oocyte sizes in the 24 hour stress duration are same data as in Fig. 2C. Size of cellularized oocytes measured in wild type (a), *gla-3(op216)* (b), and *cpb-3(op234)* (c) mutants in n= 14 to 16 worms per temperature per genotype per stress duration. d) Percentage of worms with normal oocytes (black), 1 oocyte at front (yellow), displaced oocytes (orchid), or no oocytes (white) in the 18-hour condition using the same data as in (a-c). e-h) Representative images of *gla-3(op216)* worms at 26°C for each of the oocyte patterns. (e) Normal, (f) 1 oocyte at front, (g) displaced oocytes, (h) no oocytes. Oocyte(s) encircled in orange dashed line. Box plots represent 5-95% with center line at the median and dots representing individual measurements outside the 5-95% range. *p≤0.05 significantly different within genotype between temperatures. #p ≤0.05 significantly different compared to wild type at same temperature using 2-way ANOVA with Sidak correction (a-c) or Fisher’s Exact test (d). ns p≥0.05.

**Fig. S5: Fertility decreases in wild strains under temperature stress** Experiment was done using N2 and wild strains listed in strain table. For each experiment, hermaphrodites were either maintained continually at 20°C (blue) or upshifted to 26°C (orange) or 27°C (red) starting at the L4 stage. a) Fertility was scored in n=20 to 38 worms per strain per temperature. Wild strains are ordered from smallest to largest mean brood size at 26°C. Box plots represent 5-95% with center line at the median and dots representing individual measurements outside the 5-95% range. *p≤0.05 significantly different within genotype between temperatures using 2-way ANOVA with Tukey correction. b) fertility normalized to 20°C at 26°C (orange) and 27°C (red). Dashed line represents normalized level at 20°C.

**Fig. S6: Measurements of apoptosis in wild strains under temperature stress** Experiments were done using N2 and wild strains listed in strain table. For each experiment, hermaphrodites were either maintained continually at 20°C (blue) or upshifted to 26°C (orange) or 27°C (red) for 24 hours starting at the L4 stage. Wild strains are ordered from smallest to largest mean brood size at 26°C. a) Apoptosis was measured using the *ced-1::gfp* transgene in n= 15 to 24 worms per strain per temperature. b) Apoptosis was measured using acridine orange dye in n=15 to 17 worms per strain per temperature. c) Log_2_(normalized to 20°C corpses) of apoptosis measured using the *ced-1::gfp* transgene at 26°C (orange) and 27°C (red). d) Log_2_(normalized to 20°C AO foci) of apoptosis measured using acridine orange dye at 26°C (orange) and 27°C (red). Box plots represent 5-95% with center line at the median. *p≤0.05 significantly different within genotype between temperatures using 2-way ANOVA with Tukey correction.

**Fig. S7: Wild strains with most apoptosis have largest increase during moderate temperature stress.** Correlations were created using *ced-1::gfp* apoptosis data (Fig. 3b) at 26°C and 27°C using linear regression (a and b) or second order polynomial regression (c and d). a) Mean apoptotic nuclei vs absolute increase in apoptosis at 26°C. P=0.0044. b) Mean apoptotic nuclei vs absolute increase in apoptosis at 27°C. P=0.009. c) Mean apoptotic nuclei vs % change in mean apoptosis at 26°C. P=0.6300. d) Mean apoptotic nuclei vs % change in mean apoptosis at 27°C. P=0.7036. Strains were colored by their % change of apoptosis measured using the *ced-1::gfp* transgene at 26°C compared to 20°C. Small % change in apoptosis (gold) had an increase in apoptosis of 20% – 70%. Medium % change in apoptosis (aqua) had an increase in apoptosis of 90% - 100%. Large % change in apoptosis (magenta) had an increase in apoptosis of 150% - 300%. N2 (black).

**Fig. S8: Measurements of apoptosis using act-5::yfp and ced-1::gfp in wild strains under temperature stress** Experiment was done in the N2 background using *ced-1::gfp* and *act-5::yfp* transgenes to measure apoptotic cells. Hermaphrodites were either maintained continually at 20°C (blue) or upshifted to 26°C (orange) or 27°C (red) for 24 hours starting at the L4 stage. a) Apoptosis was measured in n=15 to 17 worms per genotype per temperature. *p≤0.05 significantly different within genotype between temperatures using 2-way ANOVA with Tukey correction. b) Log_2_(normalized to 20°C corpses) of apoptosis measured using data from (a) at 26°C (orange) or 27°C (red). Box plots represent 5-95% with center line at the median.

**Fig. S9: Oocytes are larger in wild strains during moderate temperature stress** All experiments were done using N2 and wild strains listed in strain table. For each experiment, hermaphrodites were either maintained continually at 20°C (blue) or upshifted to 26°C (orange) or 27°C (red) for 24 hours starting at the L4 stage. Wild strains are ordered from smallest to largest mean brood size at 26°C. a) Number of cellularized oocytes were counted in one arm of the germline in n=14 to 18 worms per strain per temperature. b) Using the same worms as in (a), size of cellularized oocytes were measured in one arm of the germline in n=85 to 155 oocytes across 14 to 18 worms per strain per temperature. c) Size of cellularized oocytes in (b) normalized to 20°C at 26°C (orange) or 27°C (red). Dashed line represents normalized level at 20°C. d) Size of last (pink) and first (blue) cellularized oocytes from (b). Box plots represent 5-95% with center line at the median and dots representing individual measurements outside the 5-95% range. *p≤0.05 significantly different within genotype between temperatures using 2-way ANOVA with Tukey correction (a and b) or Sidak correction (d).

**Fig. S10: Embryo survival decreases and changes in L1 length are inconsistent under temperature stress.** All experiments were done using N2 and wild strains listed in strain table. For each experiment, hermaphrodites were either maintained continually at 20°C (blue) or upshifted to 26°C (orange) or 27°C (red) for 24 hours starting at the L4 stage. Wild strains are ordered from smallest to largest mean brood size at 26°C. a) Percentage of embryo survival was scored n=20 to 22 worms per strain per temperature. b) Percentage of hatched embryos normalized to 20°C at 26°C (orange) or 27°C (red). c) Length of L1 worms measured in n=35 to 75 L1s across 6 to 17 adults per strain per temperature. d) length of L1 worms normalized to 20°C at 26°C (orange) or 27°C (red). Box plots represent 5-95% with center line at the median and dots representing individual measurements outside the 5-95% range. Dashed line represents normalized level at 20°C. *p≤0.05 significantly different within genotype between temperatures using 2-way ANOVA with Tukey correction.

**Fig. S11: Apoptosis weakly relates to progeny fitness.** Correlations were created using *ced-1::gfp* apoptosis data (Fig. 3), brood size (Fig. 3), oocyte size (Fig. 4b), embryo survival (Fig. 5a), and length of L1s (Fig. 5b) at 20°C, 26°C and 27°C with linear regression (a, b, c, d, e, f, g, h) or second order polynomial regression (i, j, k, l). *p≤0.05. a) Mean brood size vs mean number of apoptotic nuclei at 20°C. P=0.3555. b) Mean oocyte size vs mean number of apoptotic nuclei at 20°C. P=0.5375. c) Mean percentage of embryo survival vs mean number of apoptotic nuclei at 20°C. P=0.8511. d) Mean length of L1s vs mean number of apoptotic nuclei at 20°C. P=0.5003. e) Mean percentage of embryo survival vs mean number of apoptotic at 26°C. P=0.4175. f) Mean length of L1s vs mean number of apoptotic nuclei at 26°C. P=0.2070. g) Mean percentage of embryo survival vs mean apoptotic nuclei at 27°C. P=0.4509. h) Mean length of L1s vs mean apoptotic nuclei at 27°C. P=0.3065. i) Mean percentage of embryo survival vs % change in mean apoptotic nuclei at 26°C. P=0.1580. j) Mean length of L1s vs % change in mean apoptotic nuclei at 26°C. P=0.4682. k) Mean percentage of embryo survival vs % change in mean apoptotic nuclei at 27°C. P=0.3110. l) Mean Length of L1s vs % change in mean apoptotic nuclei at 27°C. P=0.8442.

